## Supplementary Information for "Rational Design of Aptamer Switches with Programmable pH Response"

Ian A.P. Thompson et al.

#### Contents

**Supplementary Figure S1 |** Three-state population shift model of aptamer binding and predicted effects of construct design on binding curves
**Supplementary Figure S2 |** Three-state binding model for triplex constructs
**Supplementary Figure S3 |** Normalized pH-dependent binding curves for all triplex constructs
**Supplementary Figure S4 |** Three-state binding model for DS mismatch constructs
**Supplementary Figure S5 |** Normalized pH-dependent binding curves for A-G mismatch construct
**Supplementary Figure S6 |** Normalized pH-dependent binding curves for a construct combining triplex and mismatch control elements
**Supplementary Figure S7 |** Secondary structure optimization results for triplex sequences

**Supplementary Table S1 |** Sequences of all constructs used in this work

### Supplementary Figures

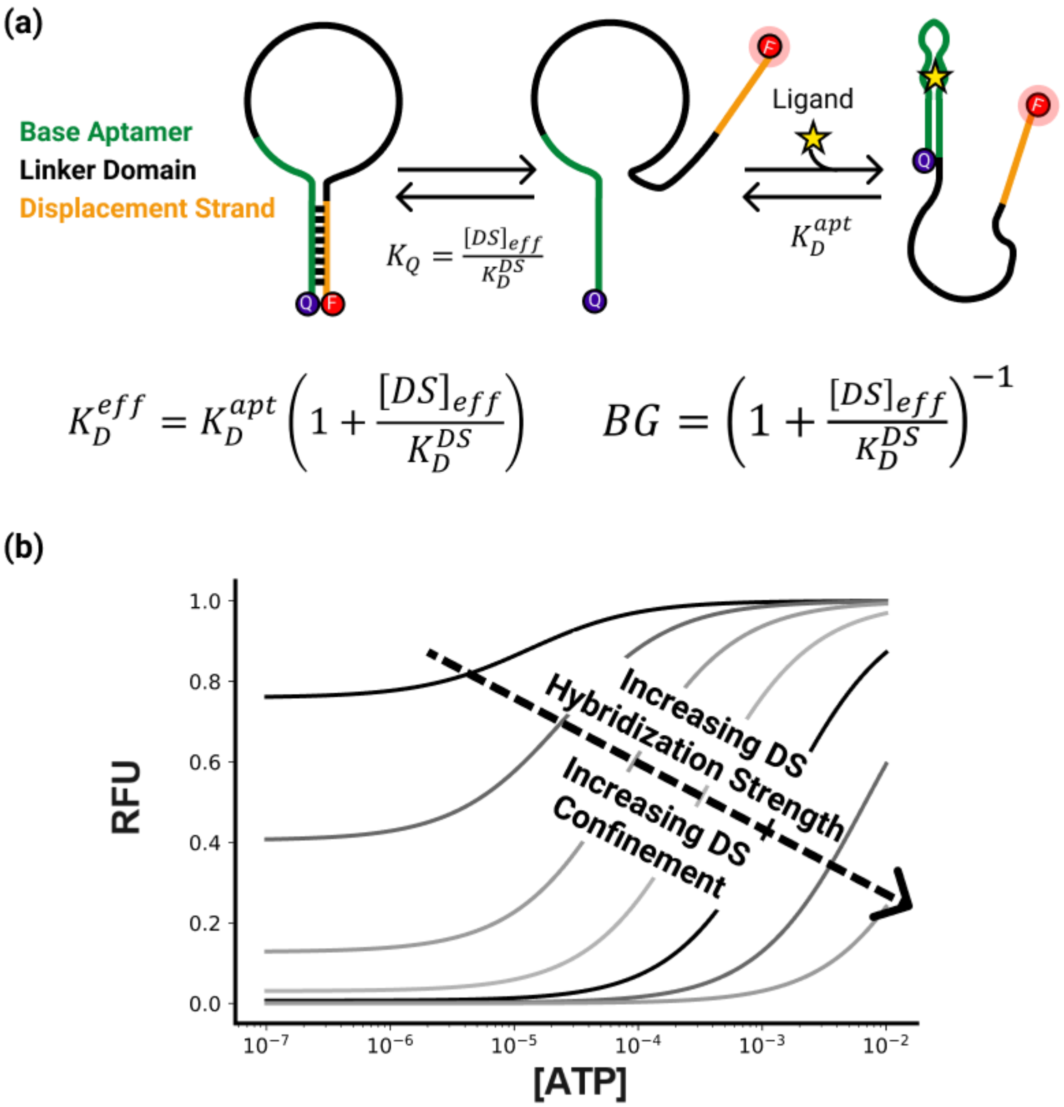


**Supplementary Figure S1 |** Three-state population-shift aptamer binding model and predicted effects of construct design on binding curves. **(a)** The three-state population shift model underlies the PSD construct mechanism, where hybridization of the displacement strand (DS) with equilibrium constant $K_Q$ inhibits aptamer binding. Target binding shifts the equilibrium away from the DS hybridized state, leading to increased fluorescence and allowing for optical quantification of target binding. The construct's effective binding affinity $K_D^{eff}$ and the magnitude of background signal both depend on the DS effective concentration, $[DS]_{eff}$, and hybridization strength, $K_D^{DS}$.[1] **(b)** Effects of varying the DS effective concentration or hybridization strength on the predicted binding curves from the three-state model.

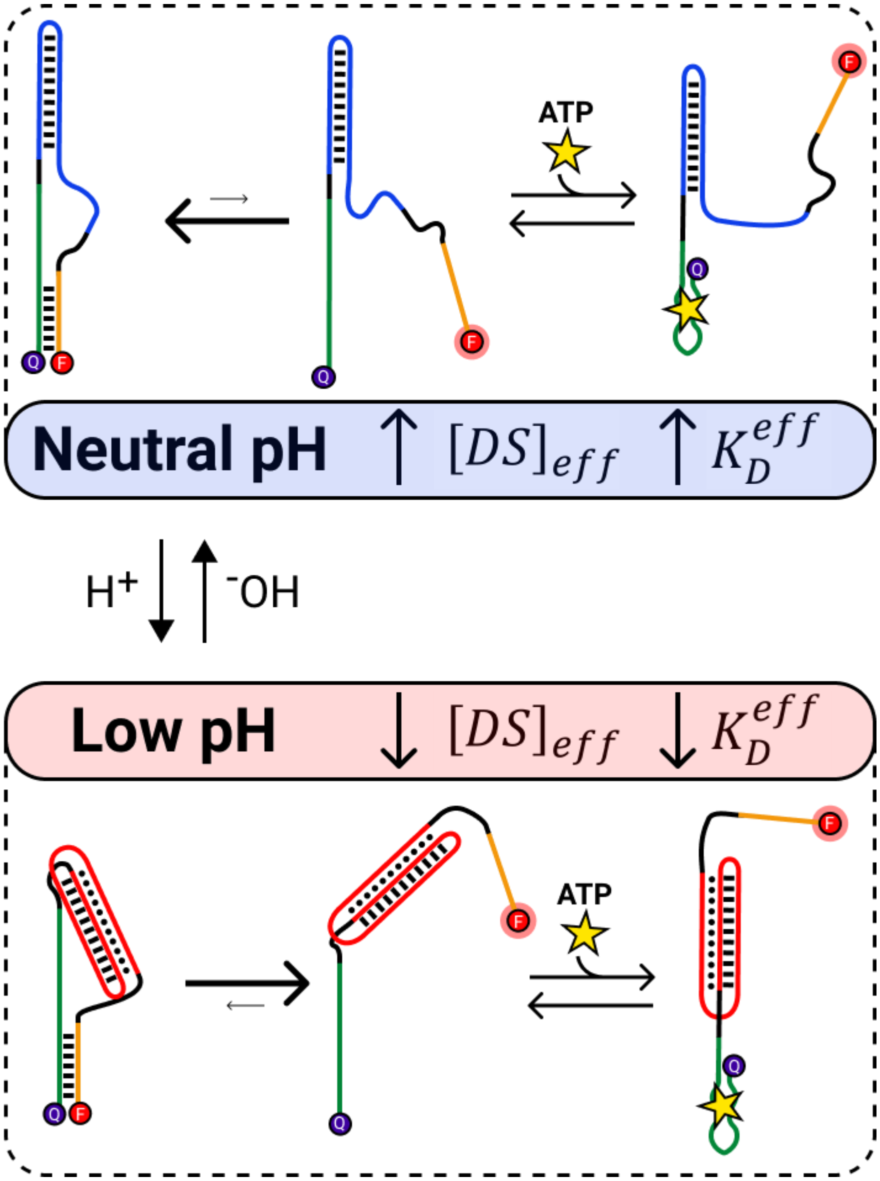


**Supplementary Figure S2 |** Three-state binding model for triplex constructs. The PSD construct mechanism relies upon pH-dependent shifts in the DS hybridization equilibrium to shift the construct's overall $K_D^{eff}$. Selective binding at low pH is achieved through pH-dependent conformational changes in the triplex motif, which lead to changes in $[DS]_{\text{eff}}$—inhibiting DS hybridization at low pH, and enhancing it at neutral pH. Equilibrium is shifted toward the hybridized state at neutral pH (top) and toward the unbound state at low pH (bottom), which selectively increases target affinity under the latter condition.

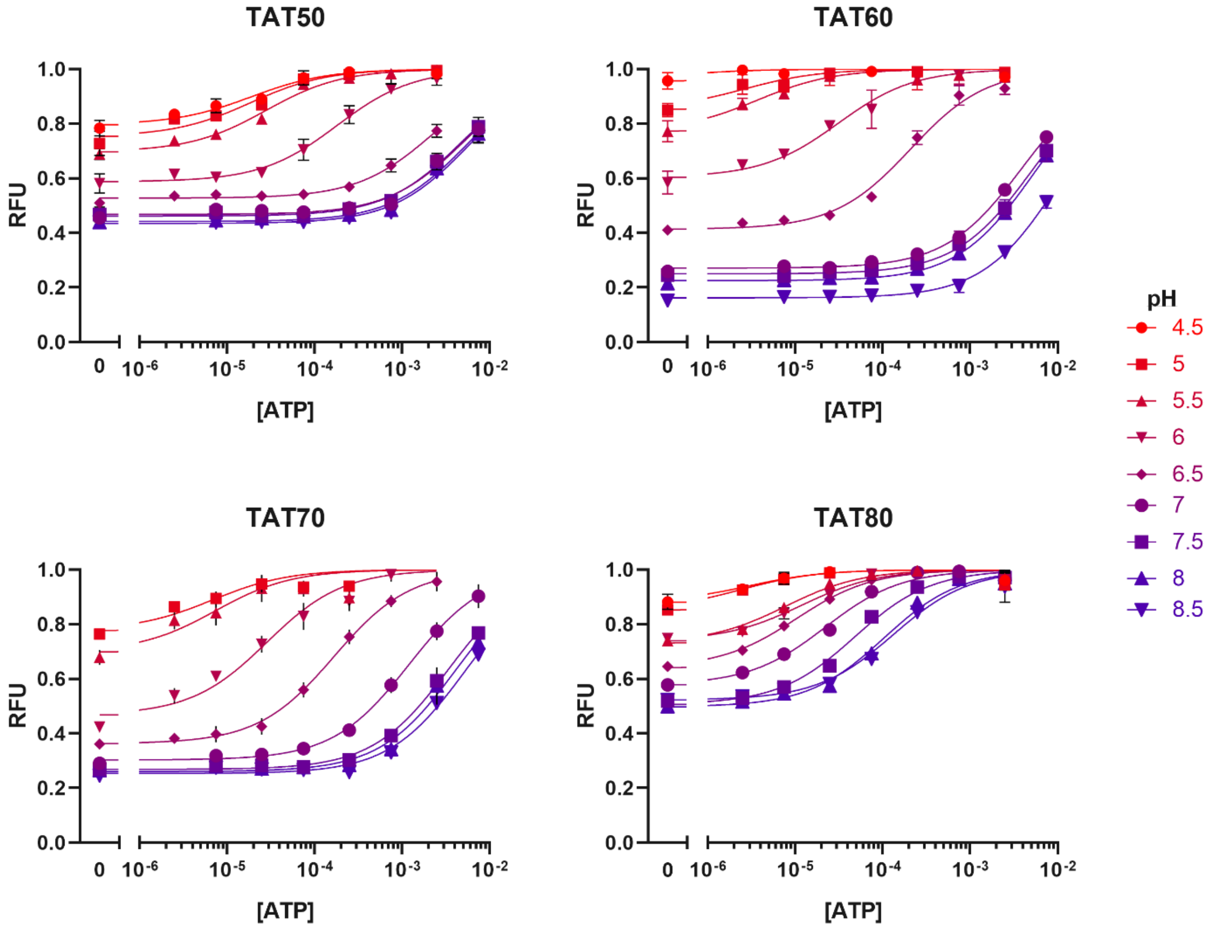


**Supplementary Figure S3 |** Normalized pH-dependent binding curves for all triplex constructs. Plots show normalized PSD construct fluorescence vs. ATP concentration at a range of solution pH values, with error bars representing the standard deviation of three replicates. The normalization procedure is given in Methods.

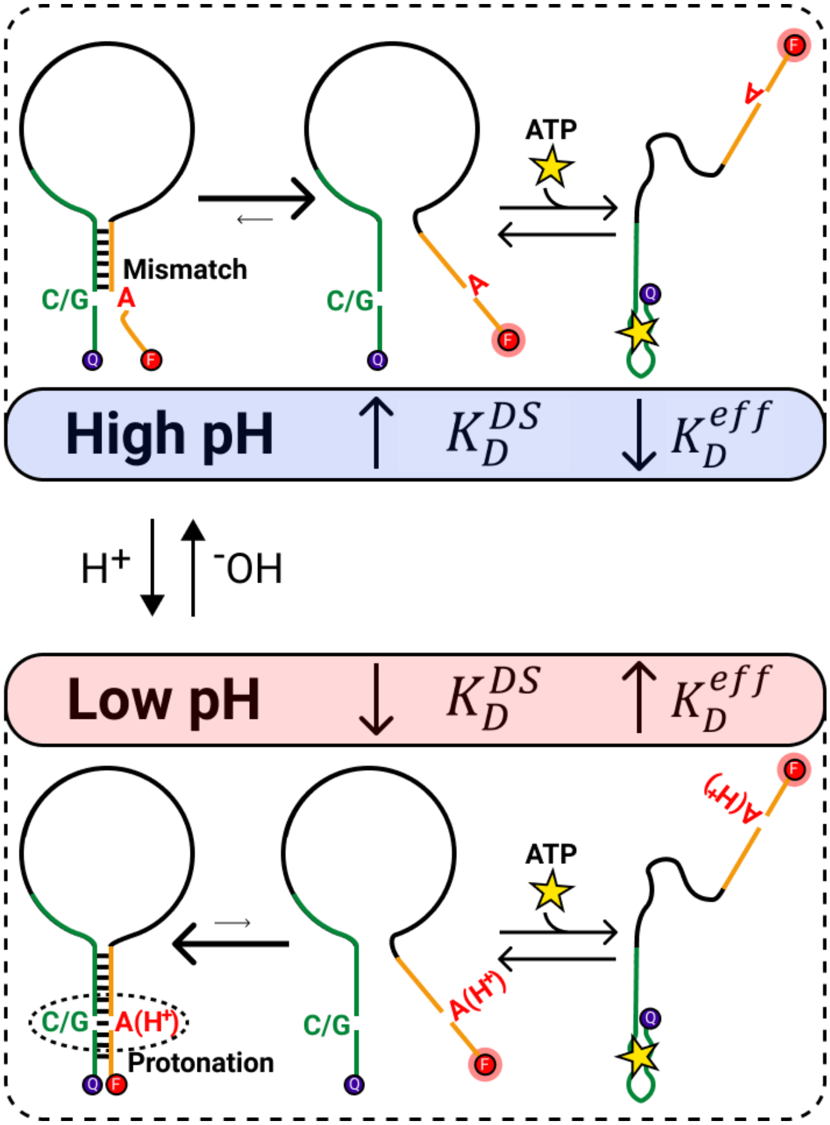


**Supplementary Figure S4 |** Three-state binding model for DS mismatch constructs. High-pH-selective binding is achieved through pH-dependent changes in DS hybridization strength due to the formation of protonated A-C or A-G mismatches. At high pH (top), the inserted mismatch remains unprotonated and does not bond, leading to weak DS hybridization (high $K_D^{DS}$) and shifting equilibrium towards the unbound state. Upon protonation at acidic pH (bottom), DS hybridization is strengthened (decreased $K_D^{DS}$) by formation of either an A(H$^+$)-C wobble base pair or an A(H$^+$)-G Hoogsteen base pair, which shifts equilibrium towards the hybridized state.

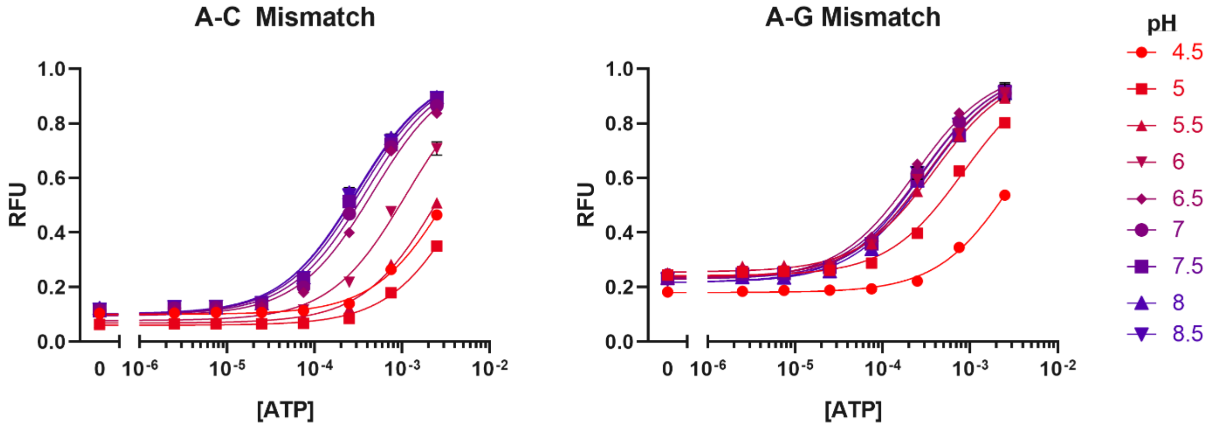


**Supplementary Figure S5 |** Normalized pH-dependent binding curves for all DS mismatch constructs. Plots show normalized PSD construct fluorescence vs. ATP concentration at a range of pH values, with error bars representing the standard deviation of three replicates. The normalization procedure is given in Methods.

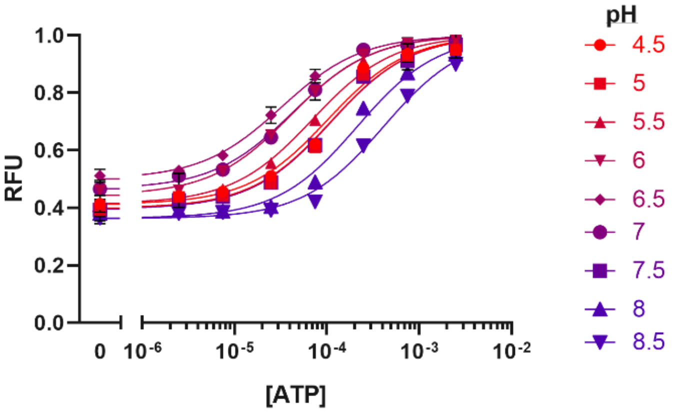


**Supplementary Figure 6 |** Normalized pH-dependent binding curves for a construct combining triplex and mismatch control elements. The tested construct incorporates both the TAT80 linker and an A-C mismatch within the displacement strand. The plot shows normalized construct fluorescence vs. ATP concentration at a range of pH values, with error bars representing the standard deviation of three replicates. The normalization procedure is given in Methods.

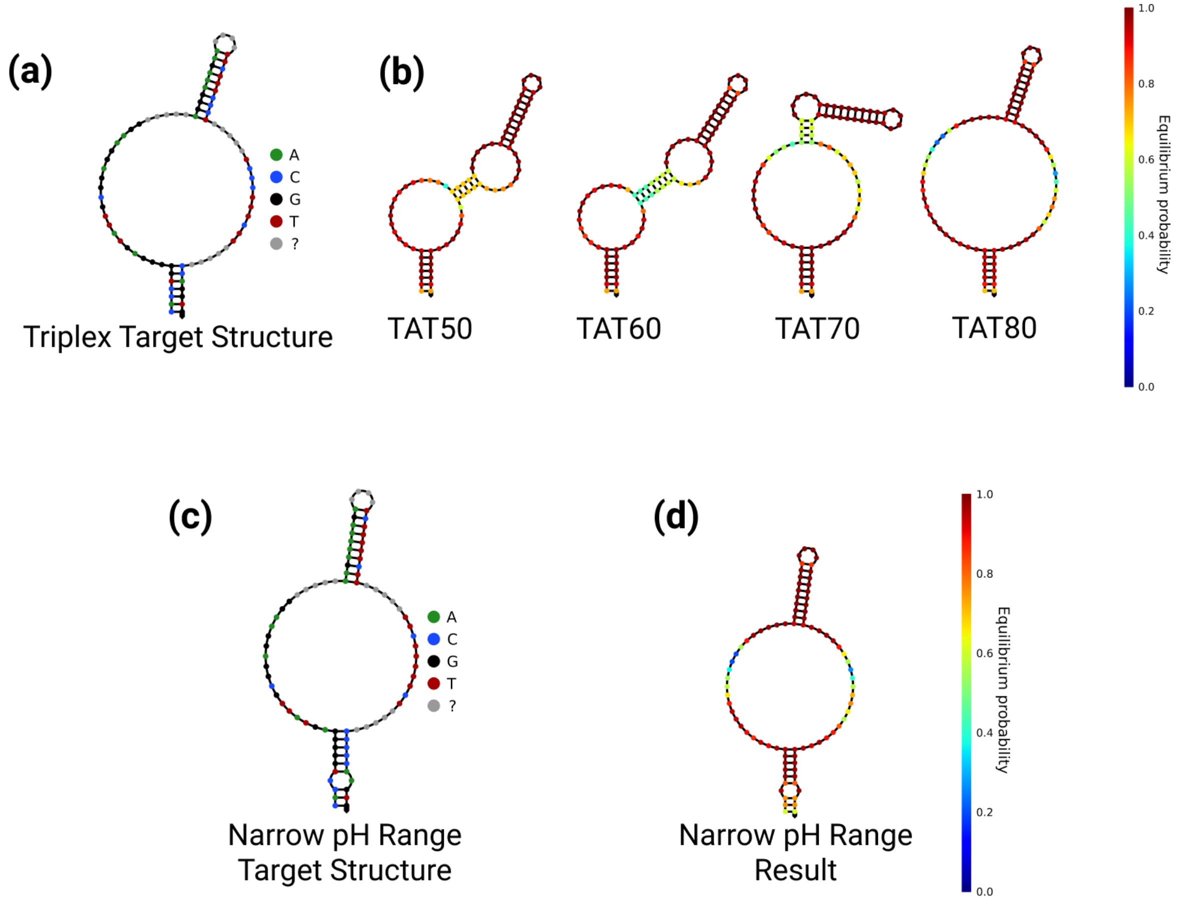


**Supplementary Figure S7 |** Secondary structure optimization results for triplex sequences. **(a)** The target secondary structure for our triplex-based PSD design was set to achieve DS hybridization to the aptamer and to allow for duplex formation between two strands of the intramolecular triplex motif. Random regions (gray bases) were optimized to minimize unintended interactions between the linker motif and aptamer. **(b)** Resulting structures of the four triplex-based PSD constructs obtained through optimization closely match the target structure, with minimized interference between the triplex motif and aptamer. **(c)** Target secondary structure and **(d)** resulting optimized secondary structure for the narrow pH range construct. All optimized structures shown were obtained through simulation in the NUPACK web application using conditions $[Na^+]$ = 50 mM and $[Mg^{2+}]$ = 6 mM at 25 °C.[2]

| Construct | | Sequence |
|---|---|---|
| ATP PSD with no triplex motif | | /5IABkFQ/CACCTGGGGGAGTATTGCGGAGGAAGGTTTTTTTTTTTTTTTTCCA GGTG/3Cy3Sp/ |
| TAT50 Triplex | Constraint | CACCTGGGGGAGTATTGCGGAGGAAGG *NNNNN* AGGGGAAGAA *NNNN* TTCTTCCCCT *NNNNN* TCCCCTTCTT *NNNNN* CCAGGTG |
| | Optimized | /5IABkFQ/CACCTGGGGGAGTATTGCGGAGGAAGG TTTTC AGGGGAAGAA GCAC TTCTTCCCCT TTTTT TCCCCTTCTT TTAAA CCAGGTG/3Cy3Sp/ |
| TAT60 Triplex | Constraint | CACCTGGGGGAGTATTGCGGAGGAAGG *NNNNN* AGGGAAAGAA *NNNN* TTCTTTCCCT *NNNNN* TCCCTTTCTT *NNNNN* CCAGGTG |
| | Optimized | /5IABkFQ/CACCTGGGGGAGTATTGCGGAGGAAGG TTTTG AGGGAAAGAA TCAT TTCTTTCCCT ATGTT TCCCTTTCTT TTTAA CCAGGTG/3Cy3Sp/ |
| TAT70 Triplex | Constraint | CACCTGGGGGAGTATTGCGGAGGAAGG *NNNNN* AGAAAGAAGA *NNNN* TCTTCTTTCT *NNNNN* TCTTTCTTCT *NNNNN* CCAGGTG |
| | Optimized | /5IABkFQ/CACCTGGGGGAGTATTGCGGAGGAAGG TTTTA AGAAAGAAGA GTGC TCTTCTTTCT CCTTT TCTTTCTTCT TTGAG CCAGGTG/3Cy3Sp/ |
| TAT80 Triplex | Constraint | CACCTGGGGGAGTATTGCGGAGGAAGG *NNNNN* AAGAAAAAGA *NNNN* TCTTTTTCTT *NNNNN* TTCTTTTTCT *NNNNN* CCAGGTG |
| | Optimized | /5IABkFQ/CACCTGGGGGAGTATTGCGGAGGAAGG TTTTA AAGAAAAAGA TTGC TCTTTTTCTT ATTGT TTCTTTTTCT AGGGG CCAGGTG/3Cy3Sp/ |
| A-C Mismatch | | /5IABkFQ/CACCTGGGGGAGTATTGCGGAGGAAGG TTTTTTTTTTTTTA CCCCCA**A**GTG/3Cy3Sp/ |
| A-G Mismatch | | /5IABkFQ/CAC CTG GGG GAG TAT TGC GGA GGA AGG TTTTTTTTTTTTTTT CCC**A**AGGTG/3Cy3Sp/ |
| Combined TAT80/A-C Mismatch | Constraint | CACCTGGGGGAGTATTGCGGAGGAAGG *NNNNN* AAGAAAAAGA *NNNN* TCTTTTTCTT *NNNNN* TTCTTTTTCT *NNNNN* CCCCCA**A**GTG |
| | Optimized | /5IABkFQ/CACCTGGGGGAGTATTGCGGAGGAAGG TTTTG AAGAAAAAGA TTGC TCTTTTTCTT ATTGT TTCTTTTTCT ATTTA CCCCCA**A**GTG/3Cy3Sp/ |

**Supplementary Table S1 |** Sequences of all constructs used in this work. Bases in italics signify unconstrained bases for optimization in Nupack. Bolded and underlined bases signify DS mismatches. /5IABkFQ/ represents the 5' Iowa Black FQ quencher modification. /3Cy3Sp/ represents the 3' Cy3 fluorophore modification.
